# High-resolution volumetric intravital imaging reveals asymmetric serotonin-dependent post-shock activity in the *Drosophila* brain

**DOI:** 10.64898/2026.05.27.728105

**Authors:** Heng Chang, Yueh-Feng Wu, Ching-Che Charng, Kai-Chun Jhan, Shun-Chi Wu, Ann-Shyn Chiang, Hsiao-Chun Amy Lin, Chung-Chuan Lo, Bi-Chang Chen, Sung-Jan Lin, Li-An Chu

## Abstract

Volumetric intravital imaging of adult *Drosophila* brains remains challenging due to optical scattering and speed limitations of conventional microscopy. Here, we present V-shape Bessel-beam light-sheet microscopy (vSPIM), an upright, high-speed platform achieving subcellular resolution across large, opaque volumes in vivo. Through real-time calcium imaging in adult *Drosophila*, vSPIM uncovered two unprecedented phenomena: first, a complex ensemble of approximately 150 olfactory projection neuron boutons (more than 400 per hemisphere) resolved 18 distinct odor valence coding patterns within the dense mushroom body calyx, aligning with connectomic architecture. Second, vSPIM revealed a previously uncharacterized asymmetric, serotonin-dependent post-shock firing (PsF) response. This PsF activity emerged within the mushroom body and dopaminergic neurons only after repetitive aversive stimulation, distinct from canonical dopamine-driven reinforcement signals. Finally, we demonstrated the versatility of vSPIM by tracking corneal endothelial dynamics during mouse wound healing. Overall, vSPIM establishes a scalable framework for intravital imaging, bridging the gap between optical innovation and circuit physiology in adult tissues.

## Introduction

Live imaging fills the temporal blind spots that limit our understanding of neural dynamics[1–3]. In particular, live volumetric imaging plays a critical role by capturing three-dimensional spatial information over time, allowing researchers to visualize neural circuit activity across both space and time. While techniques for imaging transparent models like zebrafish or *Drosophila* embryos are well-established, functional volumetric imaging of opaque, adult animals remains an unresolved limitation in the field. Adult *Drosophila* and mice offer powerful genetic tools and complex behaviors, yet their non-transparent brains scatter light, limiting the efficacy of conventional modalities. Deep-tissue methods such as multiphoton microscopy rely on point-scanning, which inherently limits temporal resolution. Although light field microscopy and spinning-disk confocal microscopy have improved imaging speeds, they often compromise imaging depth or suffer from rapid photobleaching[4, 5]. Consequently, capturing spatially distributed, sub-second neural events across large brain area remains technically demanding.

Light sheet microscopy offers an optimal balance of low phototoxicity and high spatial resolution for volumetric imaging of live specimens [6–9], it have been widely applied on real-time transparent live species such as *Drosophila* embryo and Zebrafish larvae [9]. The use of non-diffracting and self-reconstructing Bessel beams further extends the penetration depth, enabling high resolution functional imaging of deep scattering and large expanded tissues [10]. This approach has been applied to visualized intracellular dynamics and structural processes in cultured cells and live organisms such as zebrafish, *C. elegans*, and *Drosophila* larvae[11–15]. However, the feasibility of this technique in non-transparent tissues in mature living animals has not yet been well verified, especially for functional imaging within densely packed cellular structures over large areas *in vivo*.

In recent years, several advanced brain imaging technologies have been proposed to enable fast volumetric and functional recording in living animals. These milestones include high-speed whole-brain two-photon microscopy for tracking large-scale neural activity during the locomotion of behaving adult *Drosophila[16]*, as well as single-objective light-sheet microscopy optimized for resolving the sub-cellular dynamics of membranes, mitochondria, and dense-core vesicles within the living fly brain[17]. Furthermore, free-space angular-chirp-enhanced delay (FACED) microscopy introduces an all-optical, passive laser-scanning strategy that successfully bypasses the mechanical speed limitations of conventional setups, enabling ultrahigh-speed modalities such as kilohertz two-photon neural imaging[18]. Light Beads Microscopy (LBM) leverages temporal multiplexing and multi-focal axial scanning to vastly accelerate volumetric sampling rates. this framework has enabled cortex-wide volumetric imaging across multiple regions in the mouse brain at single-cell resolution, and has recently been adapted to map fly whole-brain responses evoked by individual courtship song pulses[19, 20].

Despite these notable technical milestones, capturing high-resolution dynamic neural activity across large, scattering volumes in intact mature animals remains fundamentally constrained by severe optical and biological trade-offs. To achieve high volumetric refresh rates, many existing fast-imaging platforms are forced to compromise spatial information density, relying on sparse axial scanning or heavy spatial binning. This loss of spatial resolution makes it difficult to clearly resolve fine, closely packed sub-cellular structures— such as axonal boutons within dense neuropils—where functional signals can suffer from severe spatial crosstalk. Conversely, specialized setups optimized for fine structural details frequently exhibit either heavily restricted fields of view or suboptimal volumetric update rates that fail to keep pace with rapid physiological signaling. Consequently, a critical technological gap persists in systems neuroscience: the challenge to simultaneously deliver high-density, sub-micron spatial sampling, expansive whole-brain coverage, and real-time volumetric refresh rates under low-power, biologically compatible conditions.

To overcome these persistent limitations, we developed an upright V-shape Bessel-beam light-sheet microscope (selective-plane illumination microscope, termed vSPIM), specifically engineered to bridge the gap between high-speed volumetric imaging and sub-micron structural fidelity. By exploiting the non-diffracting property of Bessel beams, vSPIM achieves a lateral resolution of ~0.7 µm and an axial resolution of ~2 µm, paired with a high-density lateral pixel sampling of 0.1625 x 0.1625 µm^2^. This architecture enables high-speed volumetric scanning of large, highly scattering tissues while preserving specimen viability, thereby eliminating the need for resolution-compromising spatial binning. Leveraging its exceptional spatial-resolving power and high signal-to-background ratio, vSPIM successfully overcomes the severe signal crosstalk typically encountered in crowded neuropils. Consequently, we resolved complex valence coding—comprising 18 distinct neural activity patterns—within the dense mushroom body calyx of the *Drosophila* brain. Furthermore, we identified a novel asymmetric, serotonin-dependent “post-shock firing” (PsF) pathway across the bilateral mushroom body, a phenomenon previously overlooked by slower/low resolution imaging modalities. This age-dependent response provides compelling evidence for prediction error encoding within the central brain. Finally, we extended the utility of vSPIM to larger mammalian tissue, successfully resolving anterior segment architecture and longitudinally tracking corneal endothelial wound healing at single-cell resolution, establishing vSPIM as a versatile and powerful tool for systems neuroscience.

## Result

### Characterization and benchmarking of the vSPIM platfrom

The system architecture and customized sample mounting framework of vSPIM are illustrated in **Fig. 1a,b** (see Methods for full architectural details). Characterization of the platform demonstrated that the self-reconstructing Bessel beam propagates up 200 µm in a transparent fluorescein solution (**Supplementary Fig. 1c**), effectively confining most excitation energy within a thin core layer and drastically reducing out-of-focus photobleaching. Point spread function (PSF) measurements using 200-nm fluorescent beads confirmed exceptional optical sectioning, yielding a sub-micron lateral resolution of 0.742 µm and an axial resolution of 2.037 µm, with a lateral pixel size of 0.1625 x 0.1625 µm^2^.

**Figure 1.**
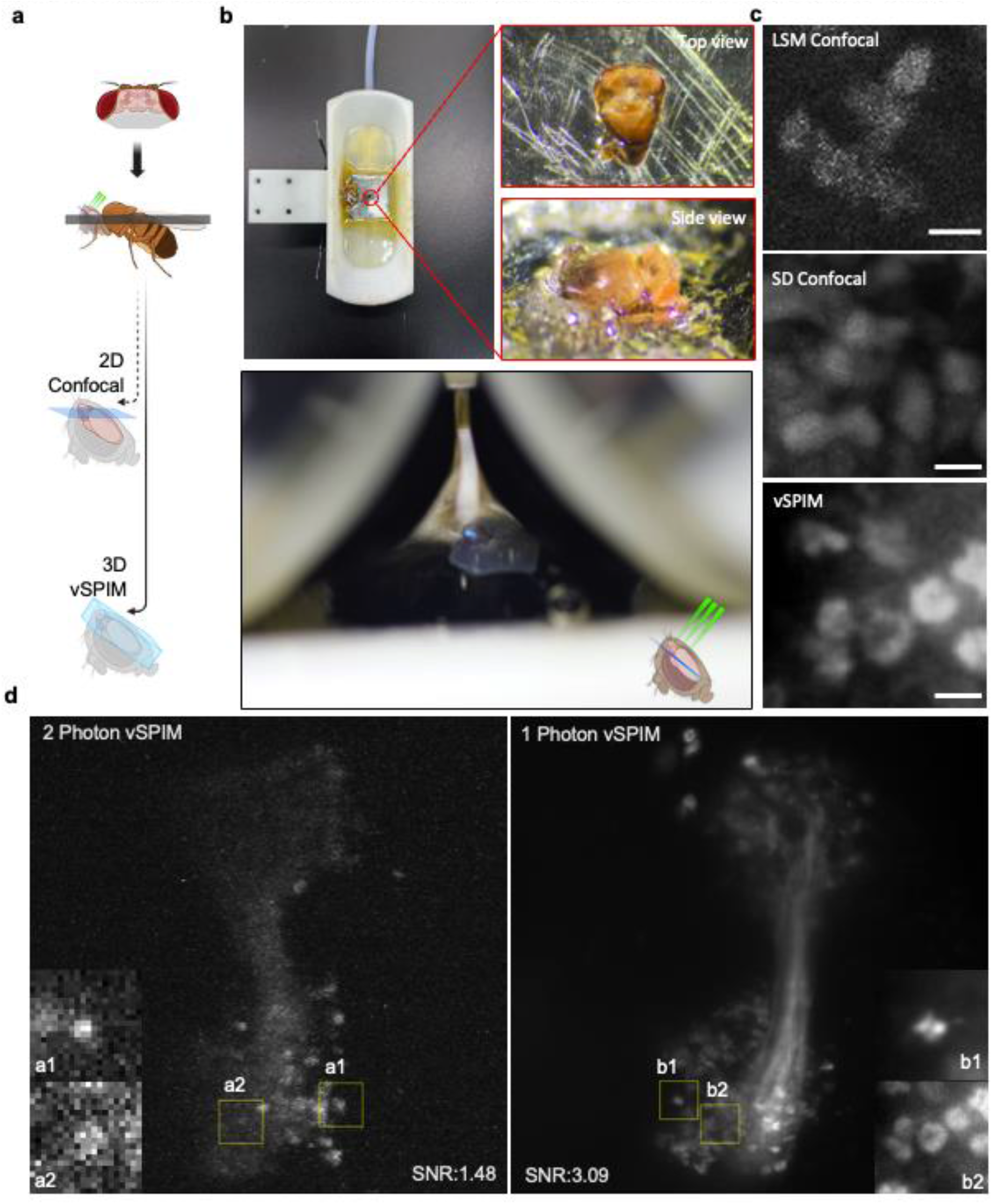
Design, sample mounting, and imaging performance characterization of vSPIM. a. Illustration of live *Drosophila* brain imaging. b. The fly is mounted secured in a custom holder. Close-up images show the fly head positioned for an upright lightsheet illumination path, along with an image of the fly during imaging. c. Representative images acquired using LSM confocal, spinning disk (SD) confocal, and vSPIM systems under identical imaging conditions, including laser power, objective magnification, numerical aperture (NA), and contrast settings. d. Comparison of 2P- and 1P-vSPIM imaging of GCaMP-labeled projection neurons under identical exposure times (70 ms). Maximum intensity projections (MIPs) are shown, with a1–b2 providing magnified single-plane views from the corresponding volumes. Compared with 2P-vSPIM (left; 4×4 binning, SNR = 1.48), which required high excitation power and did not clearly resolve densely packed boutons, 1P-vSPIM (right; 8 mW, 1×1 sampling) achieved higher contrast and resolution (SNR = 3.09). (Scale bars: 3*μm*)

Because capturing real-time spatiotemporal cellular dynamics *in vivo* demands high volumetric imaging speed, we engineered the excitation geometry and collection angle of vSPIM to enable rapid, large-scale 3D imaging of the adult *Drosophila* brain. To achieve stable intravital recording, the fly head and thorax were secured to a custom metal plate using epoxy and positioned within a dedicated holder, aligning dual water-immersion objectives at a 90° orientation (**Fig. 1b, Supplementary Fig. 1d**).

We next benchmarked vSPIM against conventional imaging modalities, including commercial laser scanning confocal microscopy (LSM) and multi-point spinning disk confocal microscopy (SD), by capturing GCaMP6m fluorescence from central brain projection neurons *GH146-Gal4 > UAS-GCaMP6m*). Under identical laser power, numerical aperture, exposure time, and contrast settings, conventional confocal systems suffered severe contrast degradation and signal attenuation in deep tissues due to optical scattering (**Fig. 1c**). Conversely, vSPIM maintained robust contrast and high structural fidelity through dense neuropils without sacrificing volumetric sampling capacity (**Fig. 1c, Supplementary Fig. 1e**).

### Challenging the 2P paradigm and cross-platform performance positioning

While prevailing literature asserts that two-photon (2P) excitation is essential for deep imaging in opaque tissues, and that 2P Bessel light-sheet microscopy should theoretically optimize contrast by eliminating sidelobe artifacts, our empirical investigations challenged this long-standing paradigm. Specifically, we evaluated imaging quality by directly comparing the 1P and 2P excitation modes of vSPIM at a fixed exposure time of 70 ms (**Fig. 1d**). Remarkably, 1P excitation achieved a significantly higher signal-to-noise ratio (SNR of 3.09 with 1×1 binning) than the 2P mode, which yielded a substandard SNR (1.48) even when driven at the maximum safe laser power threshold. To obtain usable signal in the 2P mode, 4×4 spatial binning was mandatory, which inherently degraded spatial resolution (**Fig. 1d**). Thus, despite the theoretical appeal of 2P sidelobe suppression, these counterintuitive results demonstrate that 1P Bessel excitation provides a far superior trade-off between signal fidelity, spatial resolution, and sample viability for intravital *Drosophila* imaging.

To position vSPIM within the landscape of state-of-the-art intravital functional imaging modalities, we systematically compared its core performance metrics against current beam-scanning and selective-plane illumination platforms (**Table 1**). While cutting-edge Gaussian-beam based 2P platforms, such as Light Beads Microscopy (LBM), achieve higher volume rates (~2.2Hz), they suffer from significantly coarser lateral (2.6-5µm) and axial (5-15µm) resolutions, limiting their capacity for subcellular dissection in small model organisms. Conversely, platforms offering sub-micron lateral resolution, such as LSM and FACED, are severely bottlenecked by slow volume rates (~0.167Hz) or restricted field-of-views (FOVs) and depths. Critically, vSPIM bridges this technological gap: it is the only platform that delivers an optimized balance of a ~1Hz volume rate, sub-micron lateral resolution (~0.7 µm) and an expansive voxel capacity (~71 x 10^6^ voxel), making it uniquely suited for high-throughput, high-fidelity systems neuroscience in adult *Drosophila*.

**Table 1.**
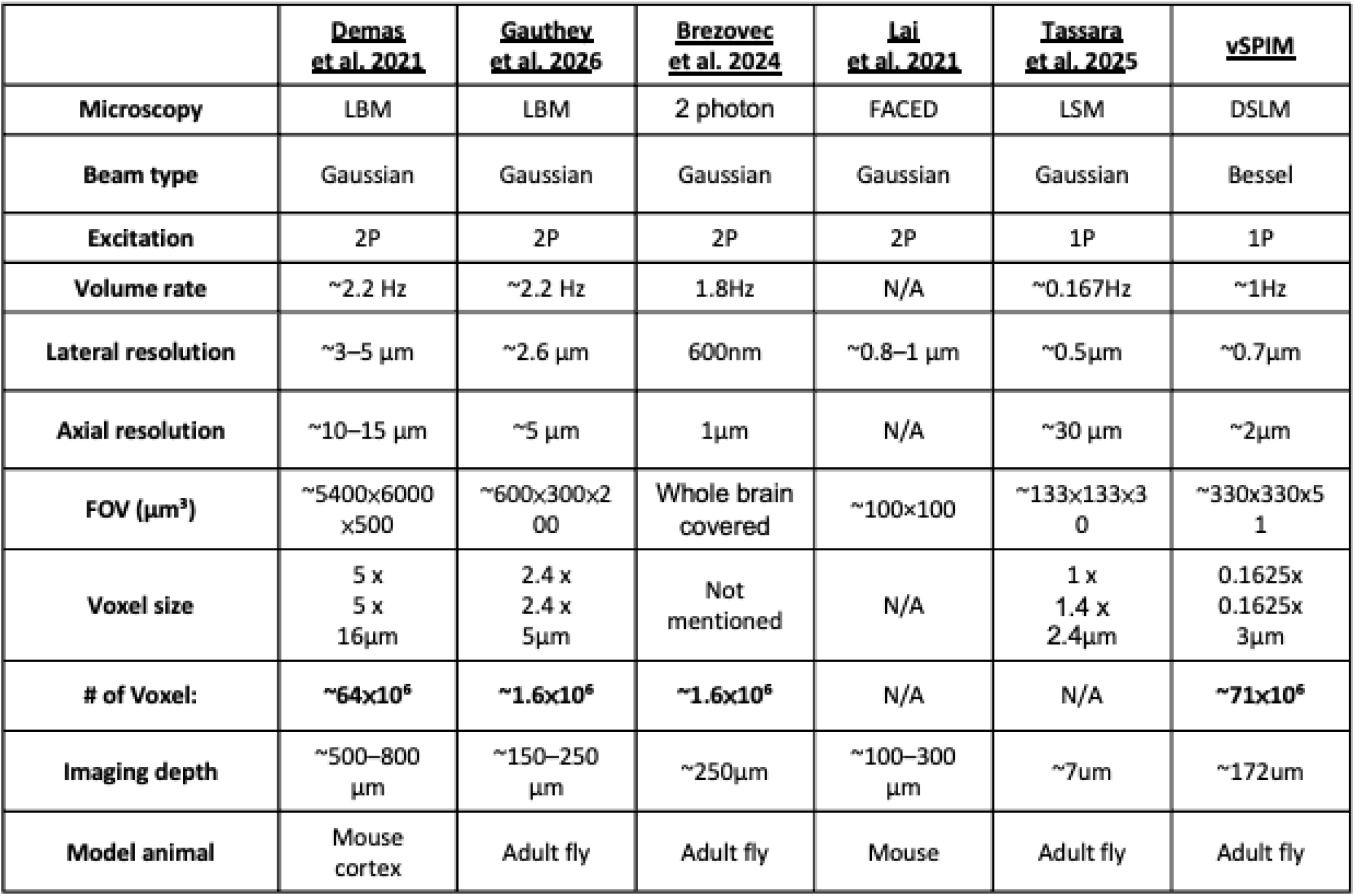
Benchmark comparison of advanced intravital microscopy systems.

### Drosophila brain calcium imaging with vSPIM

In neuroscience, tracking individual neuron activity and interactions between upstream and downstream neurons across different brain regions within the *Drosophila* entire brain has been a key focus of live imaging microscopy. In this work vSPIM offers a method to maintain high volumetric imaging speed while providing deeper illumination in fly brain. We use TH-Gal4/UAS-GCaMP6m flies to compare image quality across brain depths under three conditions: (1) cleared fly brains imaged with confocal microscopy, (2) in vivo brains imaged with confocal, and (3) in vivo brains imaged using vSPIM**(Supplementary Fig. 2a)**. As we moved deeper into the brain, confocal microscopy struggled to excite and collect fluorescence signals in living fly brain. Especially in the brain’s central regions at 73 μm depth, signal collection began to fail and worsened with increased depth. In contrast, while the lateral resolution of vSPIM at the surface was inferior to that of confocal microscopy, it maintained consistent contrast at deeper brain regions, enabling large-brain functional imaging in vivo. The structural features observed with vSPIM closely corresponded to those seen in confocal images of cleared brain tissue.

**Figure 2.**
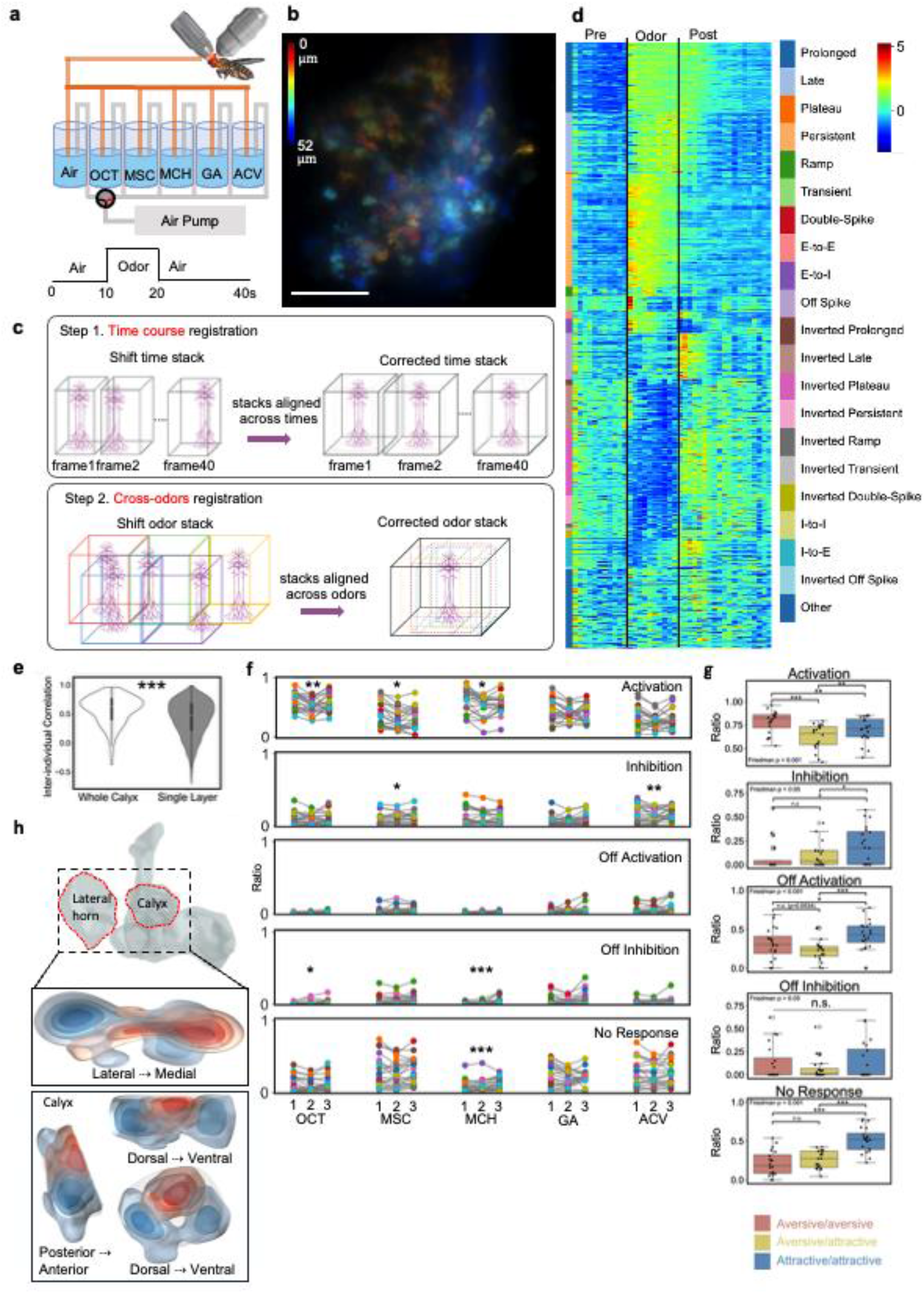
Unprecedented Functional Trace Diversity in Drosophila Olfactory Neurons. **a**. Schematic of the odor delivery setup illustrating stimulus timing and flow control during odor presentation.**b**. Volumetric vSPIM imaging of PN boutons in the calyx was performed at 1 Hz using GH146-Gal4 > GCaMP6f. **c**. A custom alignment pipeline was applied to register bouton positions across imaging depth and odor conditions. **d**. Boutons were manually annotated, and response types were classified based on calcium signal polarity and timing relative to odor stimulation. **e**. Inter-individual odor correlations were higher for whole-calyx activity than for randomly sampled single layers (two-sided Mann–Whitney U test). **f**. Depth-resolved ROI analysis revealed significant depth-dependent differences in the proportion of odor-evoked response patterns across the calyx (Friedman test followed by paired Wilcoxon signed-rank tests, FDR-corrected). **g**. ROIs exhibited significantly higher response similarity for odor pairs sharing the same valence than for pairs of opposite valences (Friedman test followed by paired Wilcoxon signed-rank tests, FDR-corrected). **h**. Spatial mapping of valence-associated responses revealed distinct depth-dependent distributions for aversive- and appetitive-preferring ROIs. *Significance thresholds: *\*p* <0.05; *\*\*p* < 0.01; *\*\*\*p* < 0.001.

Based on this result, we performed volumetric calcium imaging at 1 Hz with olfactory stimulation in *Drosophila* strains expressing GCaMP6m under the control of TH-Gal4, Pain-Gal4, and Fru-Gal4, Odor-evoked neural activity was consistently observed across multiple brain depths in different neuron types, indicating the compatibility of vSPIM to various neurological studies such as learning and memory, courtship, and pain responses **(Supplementary Fig. 2b, 2c, 2d)**.

### Quantitative odor responses and analysis in Drosophila with high resolution volumetric imaging

Olfactory information collected from the antennal lobe (AL) is transmitted via olfactory projection neurons (PNs), which bifurcate to two downstream neuropils: the lateral horn (LH) and the mushroom body (MB). In addition to examining olfactory responses across multiple brain regions, we specifically investigate how odor information is conveyed to the MB calyx through the spatially patterned axonal projections of PNs [21], using vSPIM. Within the calyx, the odor-encoding neurons—Kenyon cells (KCs)—extend multiple dendritic claws to sample inputs from PN boutons, forming discrete synaptic structures known as micro glomeruli. While the stereotyped wiring in the LH gives rise to conserved innate responses to odor stimuli, the connectivity in the MB, which supports associative learning, is markedly more complex. Recent insights into a hybrid PN-to-KC wiring scheme have emerged from the comprehensive EM-based hemibrain dataset. This structural complexity highlights the need for a complete functional characterization of odor response profiles to fully resolve the organization of olfactory projection maps in the MB—an endeavor that remains incomplete.

To validate and demonstrate the capability of our vSPIM in a densely packed neural circuit, we aimed to determine whether the full population of PN boutons within the *Drosophila* calyx exhibits valence-related spatial patterning. Odors—including two aversive (OCT and MCH), two attractive (ACV and GA), and MSC—were diluted in air and delivered toward the fly’s antennae through a custom-built odor delivery system operating at a constant flow rate of 1.25 L/min (**Fig. 2a**). The system was equipped with two-way valves to enable precise switching between odor and clean air streams. All 20 flies were exposed to an identical stimulation protocol: a 10-second air control period, followed by 10 seconds of odor presentation, then a 20-second air-infused resting phase, with 60 seconds of clean airflow between odor sequences (**Fig. 2a**).

The volumetric data are collected at a 1 Hz sampling rate (voxel size : 0.1625 x 0.1625 x 3µm^3^), covering all PN boutons within the calyx (**Fig. 2b**), by targeting *GCaMP6f* calcium sensor expression under the control of GH146-Gal4 driver line. To minimize motion artifacts affecting image analysis, we developed a dedicated alignment program that performed intra-depth alignment within each odor dataset, followed by inter-odor alignment to ensure stable bouton positions across the five odor experiments (**Fig. 2c**). Boutons were annotated manually and classified based on their calcium intensity profiles relative to odor exposure timing automatically (**Fig. 2d**). Based on ROI analysis and calcium response heatmaps, we identified up to 18 distinct neural activity patterns, including persistent activation, transient responses, and various types of inhibition, revealing an unprecedented complexity and diversity of olfactory neural responses in *Drosophila*.

Given potential variations in fixation angles or specimen biological differences during in vivo imaging, we emphasized the importance of comprehensive 3D recording. Pairwise odor correlation analyses revealed significantly higher correlation in bouton-level responses across multi-layer volume scans compared to sampled single-layer scans among 20 flies, highlighting the necessity of volumetric imaging for accurately capturing neural activity (**Fig. 2e**). To capture the potential spatial distribution differences, volumetric images of each fly were stratified into three layers. All odor response patterns were simplified into five general categories to investigate if odor distribution patterns varied across layers. The proportion of response patterns within each layer was quantified, revealing statistically significant differences across layers through Friedman testing. Specifically, MCH showed significant variations primarily in the “no response” and “off inhibition” response types across the upper, middle, and lower layers in most flies, clearly demonstrating distinct spatial distributions for odor responses (**Fig. 2f**).

Next, we further evaluated whether individual ROIs exhibited consistent response patterns across odors of the same valence, but divergent patterns for odors of opposite valence (**Fig. 2g**). The average reciprocal ratio of shared response patterns was calculated between odor pairs for each ROI. The analysis revealed that ROIs maintained significantly higher similarity within odors of the same valence and distinctly lower similarity across odors of different valences. Lastly, we highlighted the top 10% of pixels with the strongest preference for aversive over attractive odors (i.e., aversive minus attractive response) in red, and those with the strongest preference for attractive over aversive odors in blue, aggregating data across all experimental flies. The distinct spatial distribution between red and blue pixels (**Fig. 2h**) closely mirrored patterns derived from the connectome architecture[22], demonstrating strong agreement between functional fluorescent odor response imaging and structural classification, thereby reinforcing the validity of both findings.

Overall, these results comprehensively illustrate the complex and spatially specialized neural responses of the *Drosophila* calyx to odors of varying biological significance, highlighting the robustness and importance of volumetric imaging techniques such as vSPIM.

### Intravital vSPIM reveals an asymmetric, propagating post-shock firing response

Shock responses is also an important feature in fly brain for studying negative reinforcement learning and memory and pain [23]. 5-12 continuous electric shocks with a short interval between each shock are widely used as an unconditional stimulus in memory assay in *Drosophila*. We integrated the vSPIM system with an electric shock stimulation setup, targeting calcium sensor *GCaMP6f* expression in all Kenyon cells under the control of OK107-Gal4 driver. The *Drosophila* was fixed onto an iron chip and connected to a power supply following the same procedure as previous experiment (**Fig. 4a**). Throughout the experiment, the fly brain was imaged at 1 Hz with volume 330 × 330 × 154 µm^3^ (voxel size : with voxel size 0.1625 × 0.1625 × 7µm^3^) to capture the real-time response of the mushroom body (MB) to electric shock stimulation.

Upon delivering a sequence of five electrical shocks within the initial 30 seconds of training, we consistently observed robust, time-locked calcium transients in the mushroom body (MB) following each individual stimulus (**Fig. 3a,b**). Unexpectedly, vSPIM uncovered a novel neural phenomenon characterized by an extraordinarily strong calcium surge—exceeding the primary shock-evoked response amplitude by two-to five-fold—that initiated immediately after the cessation of the final shock (**Fig. 3b**). This delayed neural activity, termed post-shock firing (PsF), occurred independently of both active sensory stimulation and the initial primary calcium wave (**Supplementary Movie 2**).

**Figure 3.**
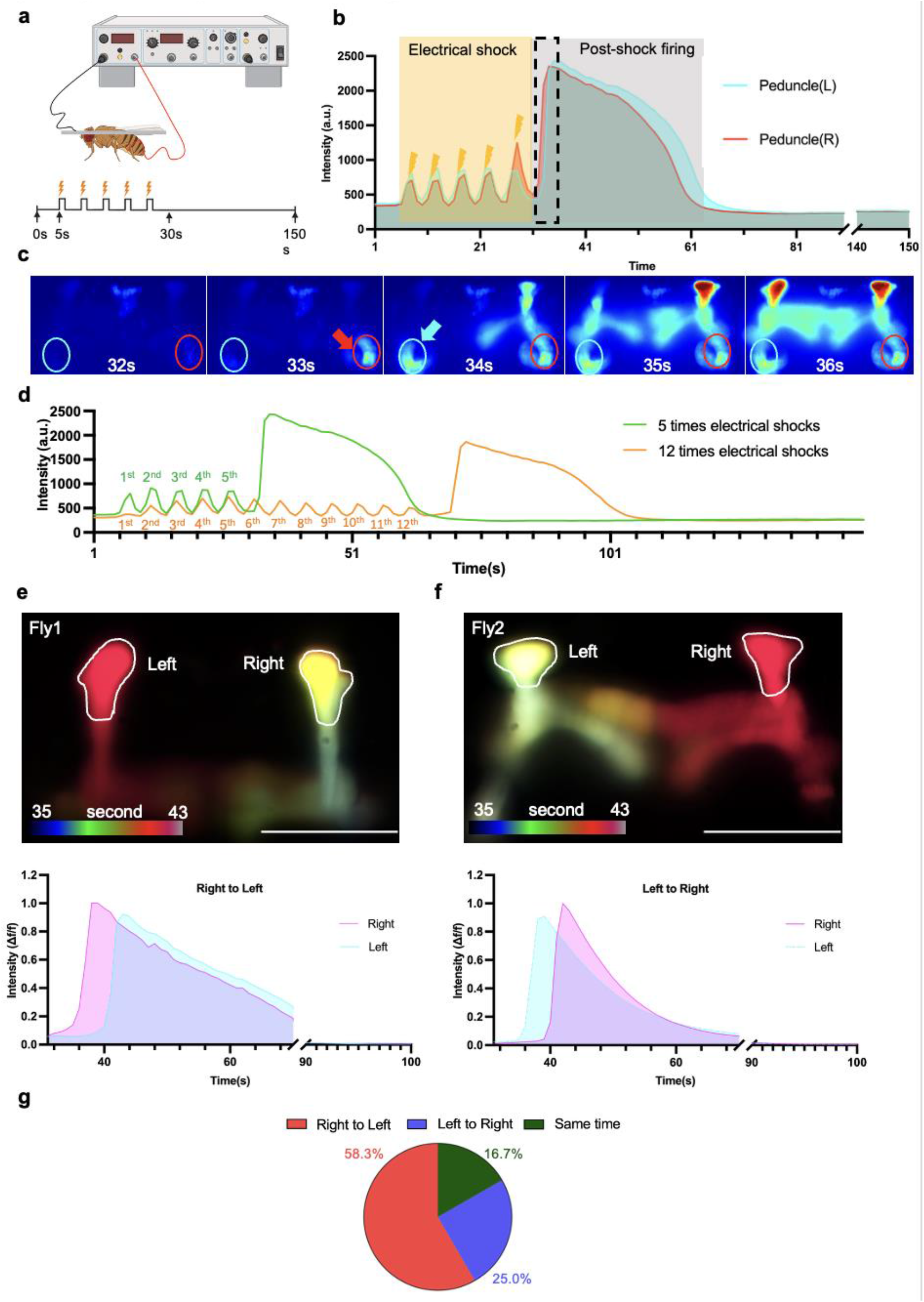
Asymmetrical prediction error response revealed with vSPIM. **a**. Schematic illustration of Drosophila electrical shock experiment. **b**. Intensity change profile of different mushroom body regions. **c**. PsF sequence in MB. Lateralized post-shock activity dynamic in the mushroom body. **d**. More electrical shocks(12 times) will hold off PsF. Color-coded time projection in **e**. R->L and **f**. L-> R flies, with Intensity profile of ROI in Alpha tips during PsF.(Scale bar, 100um). **g**. The majority exhibited right-to-left (58.3%, 7/12) propagation, suggesting a dominant lateral bias in post-shock activity flow.

Crucially, population-wide quantification across the entire MB revealed that the onset of PsF was highly asymmetrical between the left and right hemispheres, standing in stark contrast to the strictly bilaterally synchronized activity observed during the active shock period. High-resolution, frame-by-frame spatiotemporal analysis demonstrated a stereotypical directional propagation pattern of PsF. The signal typically initiated within the right gamma 1 region (red arrow) before sequentially propagating through the remaining MB lobes toward the contralateral hemisphere (**Fig. 3c, Supplementary Movie 2**). Interestingly, this lateralized initiation was not unidirectional; we observed distinct cases of both right-to-left (**Fig. 3e**) and left-to-right (**Fig. 3f**) propagation, as verified by the temporal profiles of the MB alpha tips. While individual animal variability and specific imaging orientations introduced minor variations, the overall kinetics and directional trajectories of PsF remained highly consistent. Population statistics confirmed a biased frequency, with PsF initiating predominantly from the right hemisphere (58.3%) compared to the left (25.0%) (**Fig. 3g**). To our knowledge, this represents the first visualization of a lateralized neural impulse propagating systematically across symmetrical hemispheres in the central brain.

To verify that this PsF is not merely a cumulative calcium response elicited by a 5-shock sequence, but rather a distinct predictive signal triggered specifically by the omission of an expected stimulus, we next extended the aversive training protocol from a 5-shock train over 25 seconds to a 12-shock train over 55 seconds. Intriguingly, prolonging the aversive stimulation sequence did not suppress or saturate the response. Instead, it precisely shifted the timing of the reaction, significantly delaying the temporal onset of both the resulting PsF (**Fig. 3d**) and its characteristic asymmetrical inter-hemispheric propagation (**Supplementary Fig. 3**). This time-locked emergence upon stimulus cessation strongly implies that PsF encodes a temporal prediction error, firing precisely when the brain’s expectation of an ongoing rhythmic shock is violated.

### Compartmentalized kinetics and regional selectivity of PsF within the mushroom body

To further dissect the circuit-level logic underlying this propagating response, we partitioned the MB into functional subregions and systematically analyzed their respective fluorescence intensity profiles (**Fig. 4a,b**). Our spatial decomposition revealed that distinct MB compartments exhibit markedly heterogeneous PsF kinetics and maximum intensity thresholds. Specifically, while the preceding electrical shocks elicited robust, time-locked calcium transients within the peduncle and calyx, these structural components displayed a strikingly flat, quiescent profile during the subsequent PsF phase (**Fig. 4b**).

**Figure 4.**
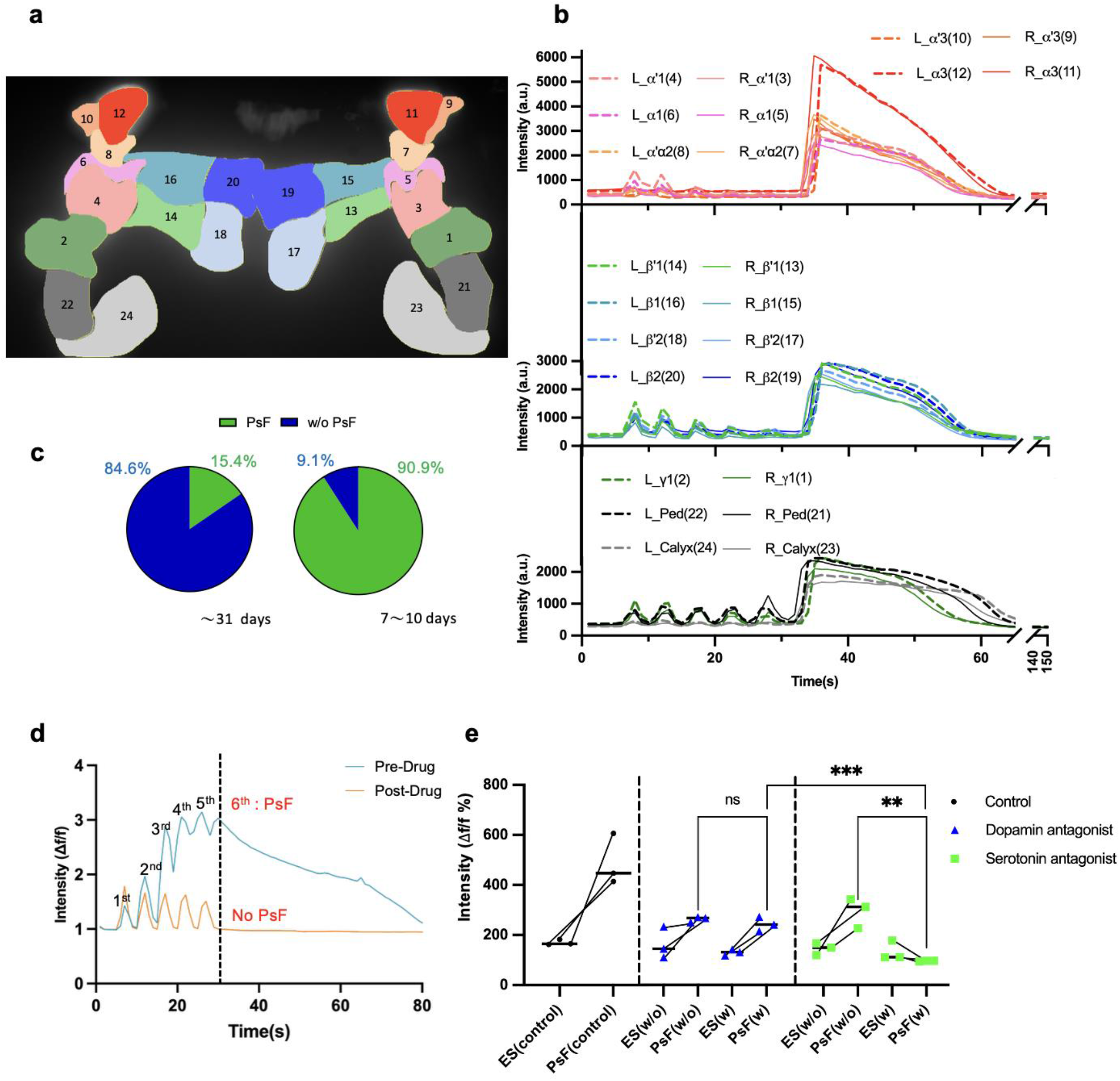
Serotonin receptor antagonist abolishes PsF in the mushroom body. **a**. Mushroom body different area sections and corresponded **b**. Intensity change profiles. **c**. Younger flies (7–10 days) displayed a significantly higher PsF occurrence (90.9%, 10/11) compared to older flies (~31 days, 15.4%, 2/13), suggesting age-related decline in PsF success rate. **d**. Normalized fluorescence intensity traces of neural activity in the serotonin antagonist group, showing strong PsF before drug administration (blue) and abolished PsF after drug treatment (orange). **e**. Serotonin receptor antagonist significantly reduced post-shock firing intensity compared to both untreated flies and those treated with dopamine receptor antagonist.

Conversely, the alpha tips exhibited the exact opposite functional profile: they remained largely unresponsive to individual electrical shocks, yet mounted an exceptionally powerful, prolonged calcium surge during the PsF event (**Fig. 4b**). This striking functional uncoupling between sensory-evoked responses and post-stimulus firing across distinct sub-neuronal compartments underscores a highly organized, regionalized processing of prediction-error-like signals within the MB network.

### Serotonergic, but not canonical dopaminergic, signaling gates PsF propagation

Previous literature establishes that nociceptive shock signals are primarily processed by peripheral pain-sensing neurons and subsequently relayed to central dopaminergic and serotonergic systems [24, 25] hat heavily innervate the MB. Consistently, we captured distinct PsF responses within dopaminergic neurons using *TH-Gal4* driven calcium imaging (**Supplementary Movie 3**). To systematically dissect the specific monoaminergic pathways driving this prediction-error-like signal, we performed targeted pharmacological blocks using flupentixol dihydrochloride (a dopamine D1/D2 receptor antagonist) and methysergide maleate (a serotonin 5-HT1/5-HT2 receptor antagonist).

Remarkably, application of the serotonin receptor antagonist completely abolished PsF propagation within the MB, while leaving the preceding sensory-evoked shock responses entirely intact (**Fig. 4e,b**). Conversely, blocking dopamine receptors neither suppressed the PsF response nor altered the primary shock-evoked calcium transients (**Fig. 4f**). This striking insensitivity to dopaminergic blockade directly challenges the classical paradigm regarding aversive reinforcement learning, where dopamine is universally recognized as the primary transmitter conveying negative valence signals to the MB [26].

While the vast majority of young adult flies (7–10 days old) exhibited robust PsF (90.9%), this neural phenomenon was largely abolished in aged flies (~31 days old), with an occurrence rate dropping to just 15.4% (**Fig. 4c**). This finding provides a critical mechanistic explanation for our age-dependent data; given that endogenous serotonin levels are documented to progressively decline in the aging *Drosophila* brain, the loss of this essential neuromodulator likely gates the observed attrition of the PsF response in aged individuals. Taken together, the prediction-error-like nature of PsF, coupled with its exclusive reliance on a serotonergic rather than a canonical dopaminergic pathway, introduces a novel neuromodulatory framework for post-stimulus circuit dynamics in the central brain.

### Intravital volumetric imaging of innervation and cellular dynamic in the mammalian anterior eye segment

Building on our success in functional imaging in *Drosophila* brain, we explored the feasibility of applying vSPIM to adult mice. The mouse anterior eye segment features an optically transparent structure with a refractive index close to that of water, making it ideally suited for water-immersion light sheet imaging[27]. However, its curved geometry (~3 mm diameter, ~1.3 mm anterior curvature[28]) poses challenges for conventional point-scanning systems, such as two-photon microscopy, which are limited by a narrow field of view and slower volumetric acquisition. In contrast, vSPIM enables high-speed, large-scale volumetric imaging with minimal phototoxicity, offering a promising solution for intact corneal imaging **(Supplementary Fig. 4)**. Consequently, we utilized mouse anterior segments, including the cornea, iris and lens, as a model to validate vSPIM for longitudinal intravital imaging of adult mouse tissues during physiological states and wound healing. To adapt the system for the anterior segment, we modified our previously developed mouse ocular surface chamber, originally designed for two-photon microscopy, for use with vSPIM via tilted objectives **(Fig.1a)**. Furthermore, we utilized five fluorescent mouse reporter strains-*GR*[29]; *mT/mG*[30], *Thy1-YFP-H*[31]; *nT/nG*[32], and αSMA^creERT2^ [33]; *tdTomato*[34]-to visualize distinct cellular and structural components in the anterior segment, including cell nuclei, cell membranes, sensory and motor nerves, and smooth muscle cells, respectively.

In the cornea, vSPIM successfully resolved three distinct cellular layers—the outer epithelium, central stroma, and inner endothelium—while preserving the native curvature and spatial integrity (Fig.5a,5b, Supplementary Movie 4). In *Thy1-YFP-H* mice, nerve fiber bundles, fine plexuses, and nodal structures were also visualized with high clarity, revealing complex innervation patterns **(Fig.5c, 5d, Supplementary Movie 6)**. Beyond the cornea, intravital imaging of the intraocular lens revealed the monolayered epithelial cells overlaying lens fiber cells, clearly delineating suture lines and the acellular lens core structure[35], consistent with known lens differentiation processes **(Fig.5e, Supplementary Movie 7–8)**. Finally, by tilting the mouse head, we resolved the circular and radial smooth muscle layers of the iris in *aSMA*^*creERT2*^; *tdTomato* mice. Real-time contraction of labeled muscle cells upon light stimulation **(Fig.5g–h, Supplementary Movie 11)** demonstrated the system’s capability for dynamic functional imaging within the intact anterior segment.

**Figure 5.**
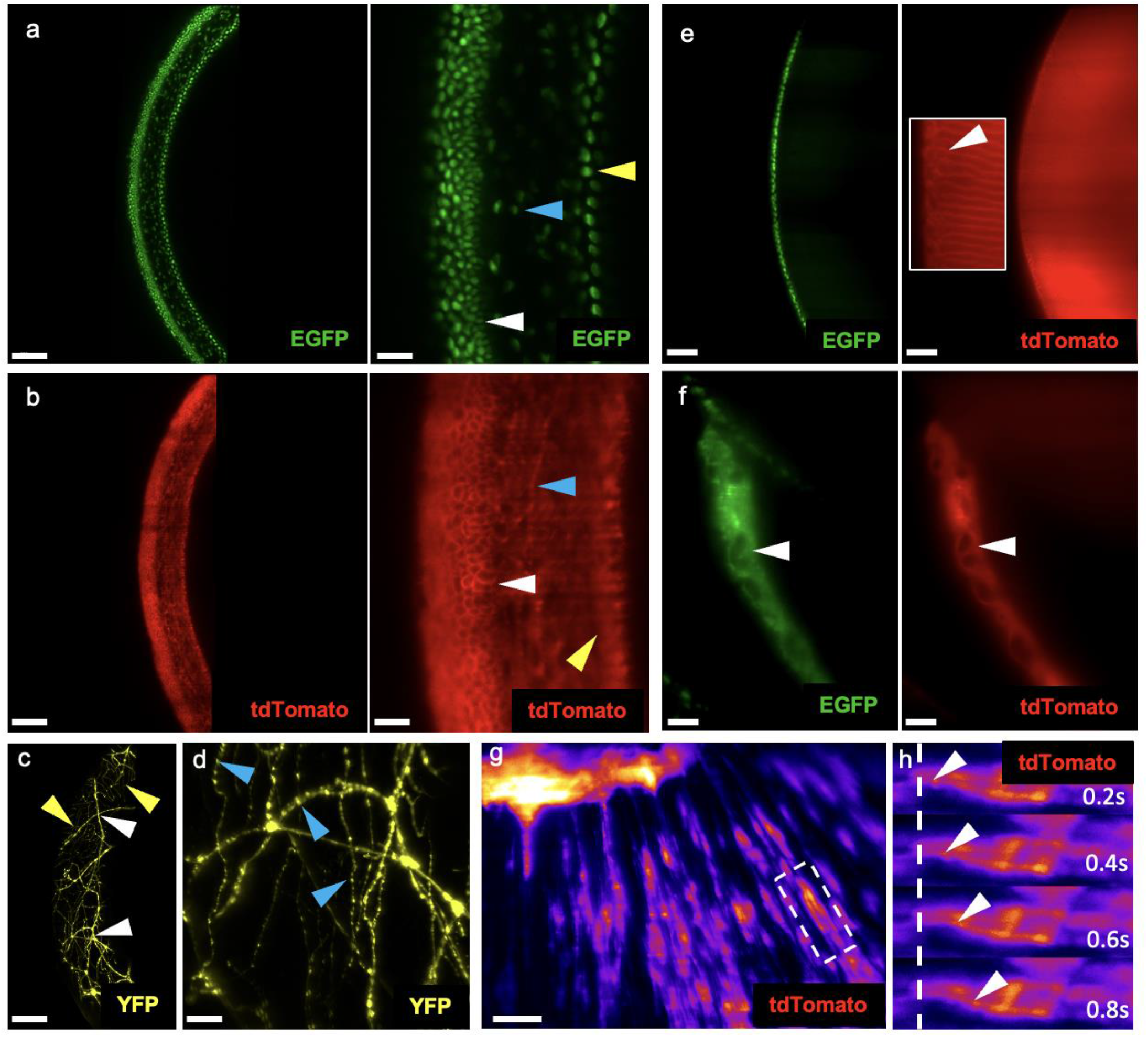
Intravital volumetric imaging of the naturally curved mouse cornea and anterior segment using vSPIM. **a,b**. Intravital 3D imaging of the cornea in *GR* (a) and *mT/mG* (b) mice. White arrows indicate epithelial cells, blue arrows indicate stromal cells, and yellow arrows indicate endothelial cells. **c,d**. Visualization of corneal innervation in *Thy1-YFP-H* mice. White arrows indicate thicker nerve bundle and branches, yellow arrows highlight fine nerve plexuses, and blue arrows mark nerve node structures. **e**. Imaging of the intraocular lens in *GR* and *mT/mG* mice. **f**. Imaging of the iris in *GR* and *mT/mG* mice. **g.h**. Iris contraction visualized in *aSMA*^*CreERT2*^; *tdTomato* mice.

### Spatiotemporal cellular dynamics of corneal endothelial wound healing following penetrating and precise injuries

Due to the limited proliferative capacity of corneal endothelial cells, wound healing in this layer primarily relies on cell enlargement and migration, resulting in a slower healing rate compared to the corneal epithelium[36–38]. While conventional histology precludes longitudinal observation of a single wound, vSPIM overcomes this limitation through its wide field of view and low phototoxicity **(Fig.5, Supplementary Fig. 4**). We first established a penetrating injury model that traverses all corneal layers **(Supplementary Figure 5)**. This injury caused a collapse of the anterior chamber and a reduction in aqueous humor space. By 24 hours post injury, the re-epithelialization was initiated rapidly; however, endothelial cells around the healing wound showed a disorganized architecture compared to the unwounded state. Fluorescent signals from keratocytes and the corneal endothelium also diminished, indicating cellular loss or potential opacity. While epithelial structure and stromal fluorescence recovered by 72-96 hours post injury, endothelial wounds remained open.

Penetrating or traumatic injuries typically induce a substantial inflammatory response, characterized by immune cell infiltration and cytokine secretion[39]. To assess healing in the absence of significant inflammation, we employed two-photon laser ablation to selectively injure the endothelium (50 μm square) without damaging the overlying epithelium or stroma **(Fig.6b, Supplementary Figure 5)**. vSPIM imaging showed that the wound remained open at 24 h but achieved near-complete closure by 48 h. The absence of observed mitotic activity supports a migration or cell enlargement-based repair mechanism. Quantification of cell density across 1000 μm^3^ in the upper, middle, and lower wound regions revealed an initial increase in density—peaking in the upper region at 24 h before returning to baseline by 48 h. Furthermore, measurements of intercellular distance (in 3-, 5-, and 9-cell clusters) showed transient expansion restricted to the upper region, consistent with active cell migration. These results confirm that while endothelial micro-wounds seal within 48 hours, the process is notably slower than epithelial healing and is driven primarily by cell motility rather than proliferation **(Supplementary Figure 6)**.

**Figure 6.**
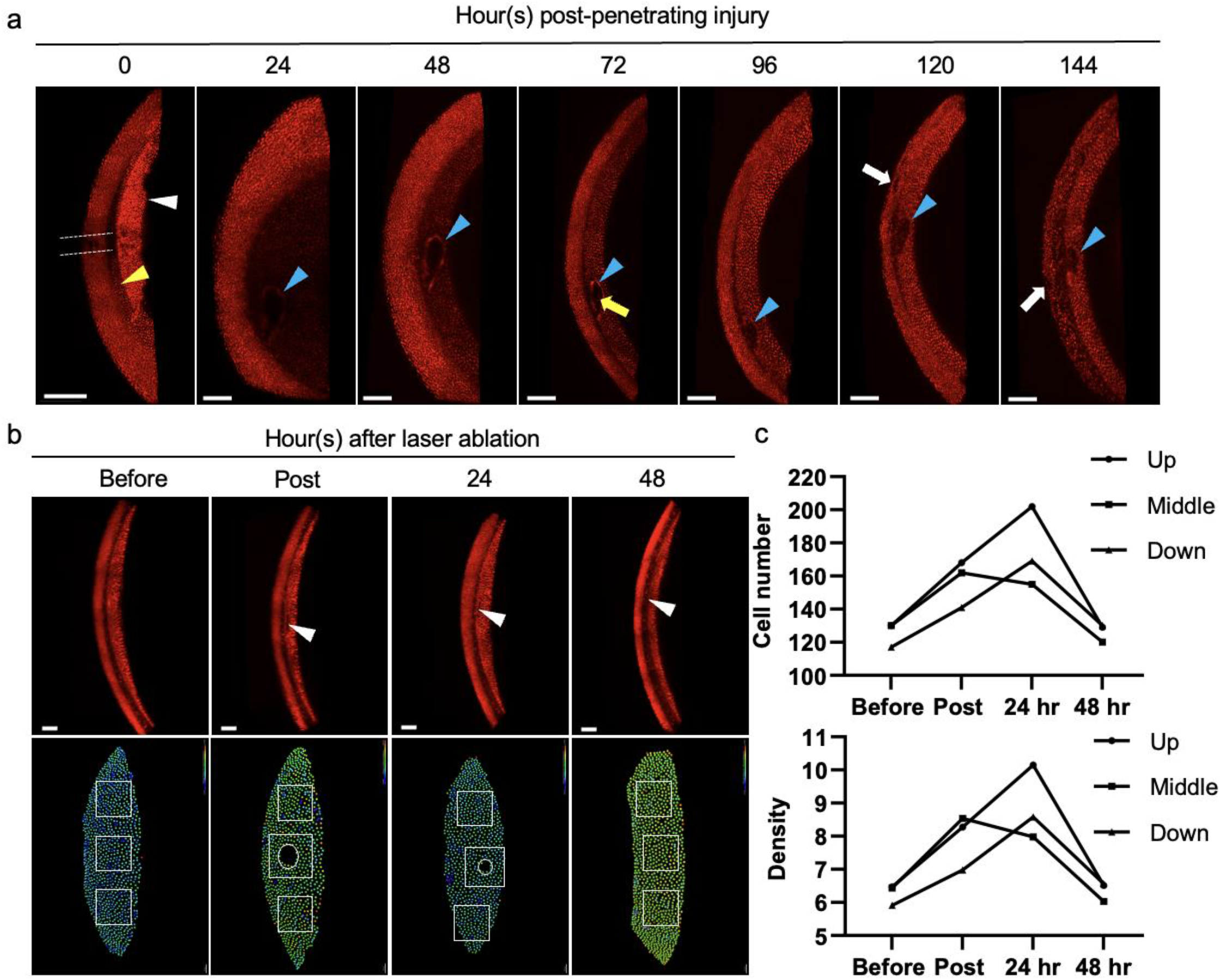
Spatiotemporal cellular dynamics of corneal endothelial wound healing visualized by vSPIM. **a**. Intravital time-lapse imaging of corneal endothelial healing following penetrating injury in *nT/nG* mice. **b**. Intravital time-lapse imaging of endothelial healing following targeted laser ablation. **c**. Quantification of cell number and density during healing derived from single-cell segmentation.

## Discussion

We introduce V-shape Bessel-beam light-sheet microscopy (vSPIM), a high-throughput, high-speed intravital imaging platform designed to resolve volumetric neural and cellular dynamics in opaque adult tissues without compromising spatial sampling. By combining a self-reconstructing Bessel beam with a unique vertical detection geometry, vSPIM achieves sub-micron lateral resolution and strong optical sectioning while minimizing phototoxicity. Crucially, vSPIM overcomes the conventional trade-off between volumetric acquisition speed and total information throughput, capturing nearly 71 million voxels per volume at a 1 Hz sampling rate. This architecture provides a 44-fold increase in spatial sampling density compared to recent high-speed whole-brain configurations (such as Light Beads Microscopy) and a 6-fold speed advantage over specialized sub-cellular light-sheet systems, while avoiding the heavy spatial binning that typically limits spatial fidelity. Furthermore, our customized specimen holder and dual-objective configuration provide the mechanical stability and clear optical access required for long-term functional recordings in complex, encapsulated organs such as the adult *Drosophila* brain. Ultimately, vSPIM bridges the gap between high-density spatial sampling and rapid temporal coverage, enabling the mapping of distributed, transient cellular activities across intact neural networks.

### Volumetric neural coding in the adult Drosophila brain

Applying vSPIM to the adult *Drosophila* brain enabled direct visualization of distributed neural activity patterns across densely packed regions at sub-second temporal resolution. The resulting volumetric datasets uncovered layer-specific odor valence coding in the mushroom body calyx (**Fig. 2**), where projection neuron boutons exhibited spatially organized response profiles consistent with connectomic predictions[40]. This observation demonstrates that volumetric sampling captures circuit-level functional diversity that planar imaging or partial-volume recordings often overlook. Moreover, alignment of odor response patterns across layers revealed that valence information is not uniformly represented but stratified, reflecting distinct synaptic integration modes within the calyx micro glomeruli.

Beyond sensory coding, vSPIM revealed a previously undescribed asymmetric, serotonin-dependent post-shock firing (PsF) in the mushroom body (**Fig.3,4**). PsF emerges only after repeated aversive stimulation, propagates unilaterally across lobes, and disappears upon serotonin receptor blockade while remaining insensitive to dopamine antagonism. This finding challenges canonical models of aversive learning in which dopaminergic neurons convey punishment prediction signals to the mushroom body[41, 42]. Instead, the delayed and serotonin-dependent nature of PsF suggests a post-event prediction-error–like signal, potentially encoding mismatch between expected and actual termination of aversive input[43], which may represent a conserved mechanism of aversive prediction updating. The pronounced age dependence of PsF further implies that serotonergic modulation may contribute to the degradation of aversive learning plasticity during aging. Together, these results underscore the utility of vSPIM for discovering emergent circuit-level phenomena that unfold over distributed networks in vivo.

### Extending vSPIM to mammalian tissues

To evaluate the versatility of vSPIM beyond invertebrate models, we adapted the platform for large-scale imaging of the mouse anterior eye segment, which features naturally curved geometries that pose challenges to conventional point-scanning microscopy (**Fig. 5)**. The upright configuration and extended depth of field afforded by the Bessel beam allowed stable, minimally invasive imaging of the intact cornea, iris, and intraocular lens under physiological conditions. Using multiple reporter strains, vSPIM resolved fine structures such as epithelial layers, sub-basal nerve plexuses, lens fiber organization, and smooth muscle dynamics during iris contraction, highlighting its compatibility with complex three-dimensional geometries.

Crucially, vSPIM enabled longitudinal tracking of corneal wound healing across three cell layers at single-cell resolution following both penetrating and precisely targeted endothelial injuries (**Fig. 6**). Our results revealed that endothelial wound healing proceeded primarily through collective cell migration and enlargement rather than proliferation, consistent with the limited regenerative capacity of the corneal endothelium. By capturing real-time repair dynamics across wide fields of view without physical sectioning and with minimal phototoxicity, vSPIM provides a new window into corneal physiology and regenerative mechanisms. These findings demonstrate that vSPIM can bridge spatial and temporal scales—from subcellular events in neural circuits to tissue-level regeneration in mammals—using a unified optical platform.

### Conceptual implications and future perspectives

Together, these results establish vSPIM as a generalizable framework for intravital volumetric imaging across biological scales. The combination of high-speed acquisition, optical sectioning depth, and mechanical stability expands the range of tissues accessible for functional imaging—from adult neural networks to regenerative epithelia. The discovery of the serotonin-dependent PsF in *Drosophila* further exemplifies how enhanced spatiotemporal coverage can uncover neuromodulatory processes that have eluded prior imaging approaches. In the mammalian system, the ability to monitor wound healing dynamics continuously and noninvasively opens new possibilities for studying ocular pathophysiology, stem-cell activity, and drug responses in vivo.

Looking forward, incorporating adaptive optics, remote focusing, and dual-view detection will further improve isotropy and penetration depth, while integration with behavioral assays and electrophysiology could link large-scale circuit dynamics with sensory processing and motor output. By uniting optical innovation with system-level neuroscience, vSPIM overcomes the trade-off between imaging speed and depth in opaque tissues. The discovery of serotonin-mediated PsF highlights the system’s potential to reveal emergent circuit properties that are undetectable with standard planar imaging. As a versatile platform, vSPIM paves the way for comprehensive 4D mapping of structure–function relationships, spanning from emergent circuit properties in nervous systems to dynamic regenerative mechanisms in complex mammalian organs.

## Methods

### vSPIM system setup and image acquisition

We started by expanding a Gaussian laser beam using a beam expander. The expanded beam was then directed onto an annular ring mask, creating a uniform ring shape. This patterned light was subsequently conjugately projected onto the rear focal plane of a commercial objective lens. After focusing on the excitation objective, we can produce Bessel beams with different lengths and thicknesses by controlling the inner and outer radius of the ring pattern on the mask. A rotating galvo mirror moves the Bessel beam horizontally across the imaging plane, creating a time-averaged virtual lightsheet at the focal plane of the excitation objective lens. The detection objective placed at a 90° orientation relative to the excitation objective, with its imaging plane aligned with the axial plane of the excitation objective. The coordinated movement of another rotating galvo mirror and a piezo stage equipped with the detection objective lens ensures that the illumination and imaging planes move confocally. These components are precisely synchronized, allowing for high-resolution volumetric images. Several relay lens pairs are used between mask and excitation objective to maintain ring light shape and power. This setup ensures the light sheet maintains its integrity and power throughout the optical path. Finally, after the detection objective lens and the tube lens, the images are acquired by a 2D camera.

The optical setup is illustrated in Fig. 1a. A 488 nm laser source (Coherent OBIS 488 LS, 150 mW) emits a beam that is first collimated to a 1/e^2^ diameter of 2.5 mm using two lenses: L1 (7.5 mm focal length, 5 mm diameter, Thorlabs AC050-008-A-ML) and L2 (30 mm focal length, 12.5 mm diameter, Thorlabs AC127-030-A). The collimated beam is directed through an acousto-optic tunable filter (AOTF; AOTFnC-400.650-TN, AA Quanta Tech, Optoelectronic), which regulates both the illumination duration and laser intensity. Following the AOTF, the beam is expanded to a 1/e^2^ diameter of 5 mm using a pair of achromatic lenses configured as a beam expander (Thorlabs AC254-125-A [L7] and AC254-250-A [L8], Ø1”, 400–750 nm). This expanded beam uniformly illuminates a custom annular mask with a 1,500 Å aluminum coating. A relay lens pair (Thorlabs AC254-125-A [L12] and AC254-100-A [L13], Ø1”, 400–750 nm) conjugates the ring pattern from the mask to the galvanometric scanner system (Cambridge Technology 6215H). The scanners are aligned in a 4f configuration using two achromatic lenses (Thorlabs AC254-100-A [L14, L15], Ø1”, 400– 750 nm). After scanning, the beam passes through another relay lens system (Thorlabs AC254-125-A [L16] and AC508-300-A [L17], Ø2”, 400–750 nm) that magnifies the ring pattern and aligns it to the back focal plane of the excitation objective (Nikon CFI Apo NIR 40XW, 0.8 NA, 3.5 mm WD). This configuration generates a self-reconstructing Bessel beam via optical interference, maintaining high energy confinement within the illumination plane while supporting extended axial propagation. Perpendicular to the illumination axis, a second water-dipping objective is mounted on a piezoelectric scanner (Physik Instrument P-725.4CD PIFOC) to collect fluorescence. The emitted signal is filtered through a quad-band emission filter (Edmund Optics #87247, 440/521/607/700 nm, 25 mm diameter) and imaged onto a scientific CMOS camera (Hamamatsu ORCA-Flash4.0 V3) via a 200 mm tube lens (Thorlabs AC254-200-A, Ø1”, 400–750 nm).

### PSF measurement

To characterize the system’s point spread function (PSF), fluorescent microspheres (Thermo Scientific, Fluoro-Max, G200B) were adhered to a Poly-L-Lysine-coated coverslip (Sigma, P4707) to ensure immobilization. The PSF was acquired by scanning the light sheet across a 20 μm axial range with a z-increment of 500 nm.

### Fly stocks

Fly stocks were maintained at 25 °C with a relative humidity of approximately 70% under standard conditions of a 12-hour light/dark cycle, using cornmeal-based food. This study employed the following fly strains: GH146-Gal4 (BDSC 30026) targeting olfactory projection neurons; Pain-Gal4 (BDSC 27894) for labeling sensory neurons involved in nociception; OK107-Gal4 (BDSC 854) for mushroom body Kenyon cell-specific expression; Fru-Gal4 (BDSC 30027) to selectively label neurons regulated by fruitless sequences; and TH-Gal4 (BDSC 8848) for marking dopaminergic neuronal populations. Additionally, UAS-GCaMP6m (BDSC 42750) and UAS-GCaMP6f (BDSC 42747) served as reporters for calcium activity, enabling the visualization of neural dynamics in cells expressing Gal4 drivers.

### Fly mounting and dissection protocol

Flies were positioned its’ head and upper chest passed through the central opening of the metal plate in supplementary figure 1, glued by using AB glue (DEVCON, 5 minutes Epoxy). The orientation of each fly was carefully adjusted to align the brain region of interest with the optical path, thereby minimizing the amount of tissue the excitation and emission light needed to pass through. To expose the brain, a rectangular window was created on the head capsule using fine scalpels, with cuts made at both the anterior and posterior sides. Surrounding fat tissue was gently displaced to the edges of the opening, and tracheal branches were severed and moved aside to provide unobstructed optical access to the brain. All dissection procedures were performed under adult hemolymph-like (AHL) saline composed of 108 mM NaCl, 5 mM trehalose, 10 mM sucrose, 2 mM KCl, 2 mM CaCl_2_·2H_2_O, 8.2 mM MgCl_2_·6H_2_O, 4 mM NaHCO3, 1 mM NaH_2_PO_4_·H_2_O, and 5 mM HEPES.

### Odor delivery system

A continuous background airflow was maintained at a rate of 1 L/min. A secondary air stream, flowing at 200 mL/min, was directed through either a vial containing mineral oil (for control conditions) or an odorant vial, depending on the position of a manually operated valve. The odorant vials contained chemical stimuli including 4-methylcyclohexanol (MCH; Sigma-Aldrich, 153095), 3-octanol (OCT; Sigma-Aldrich, 218405), Geranyl Acetate (GA; Sigma-Aldrich, 173495), Methyl Salicylate (MSC; Sigma-Aldrich, 76631), and Apple Cider Vinegar (ACV; Kong Yen, 12419). Each odorant was prepared by diluting 10 μL of the pure compound into 10 mL of mineral oil (TEDIA, MR1285-005). This setup resulted in a final odorant dilution of 1:1000 at the level of the fly antennae.

### Diversity Analysis of Odor-Evoked Calcium Responses

Functional imaging data were first cropped and warped using imageJ, then processed in two ways: (i) manually annotated bouton-level analysis, and (ii) voxel-based analysis via spatial binning along the x- and y-axes (final resolution: 3.25µm). Calcium traces were classified in two stages. First, we used rule-based criteria to assign each trace to “Activation” or “Inhibition” if the peak (|ΔF/F| > 0.1) occurred during odor stimulation, or “Off activation” / “Off inhibition” if the peak followed stimulation. Traces below threshold were labeled “No response.” Second, we matched trace dynamics to a library of prototype patterns (e.g., Transient, Ramp, Double spike, and their Inverted forms). Traces were min–max normalized and assigned by Pearson correlation (threshold: 0.7); others were labeled “Others.” We compared the number of distinct response patterns between odors of different valences using the Wilcoxon rank-sum test. OCT and MCH were categorized as aversive odors, while ACV and GA were classified as attractive odors. For trace repertoire visualization, all traces were z-score normalized and pooled across odors.

### Inter-Fly Odor Response Correlation

To assess inter-individual similarity in odor-evoked responses, we extracted calcium traces from all ROIs (boutons) and computed pairwise odor correlations using Pearson correlation across full temporal profiles. These odor-wise correlations were then used to quantify pairwise inter-fly response similarity.

### Spatial Analysis of Odor Response

To examine the spatial distribution of odor responses, ROIs were divided into three groups based on z-coordinate quantiles. Within each group, the proportion of response pattern types was quantified, and statistical differences across groups were assessed using the Friedman test. To further explore spatial patterns, we analyzed voxel-level data, which provide more continuous coverage of odor responses. To contrast odors by valence (aversive vs. attractive), voxel-wise responses to attractive odors were subtracted from those to aversive odors. The top 10% of voxels showing the strongest aversive-biased responses were highlighted in red, while the bottom 10%—those with strongest attractive-biased responses—were highlighted in blue. Kernel density estimation (KDE) was then used to estimate spatial distributions, visualized using the *mayavi* package.

### ROI Response Similarity Analysis

To assess whether individual ROIs exhibit consistent response patterns across odors of the same valence and divergent patterns across opposite valences, we computed the average reciprocal ratio of shared response patterns between odor pairs. Specifically, for each ROI and odor pair, we calculated the proportion of matching response patterns in one odor that also appeared in the other, and averaged this ratio in both directions.

### Transgenic mice for ocular anterior segment imaging

*GR* (B6;129-Gt (ROSA)26Sor^tm1Ytchn^/J, #021847), *mT/mG* (Gt(ROSA26)^ACTB-tdTomato-EGFP^, #007676), *Thy1-YFP-H* (B6.Cg-Tg(Thy1-YFP)HJrs/J, #003782), *nT/nG* (B6N.129S6-Gt(ROSA)26Sor^tm1(CAG-tdTomato*,-EGFP*)^ Ees/J, #023537) and *tdTomato* (B6.Cg-Gt(ROSA)26Sor^tm14(CAG-tdTomato)Hze^/J, #:007914)mice were from The Jackson Laboratory. The *aSMA*^*creERT2*^ mouse line was kindly provided by Prof. Pierre Chambon (Wendling, Bornert et al. 2009). *R26R-GR* mice express a nuclear H2B-EGFP and a membrane-bound mCherry-GPI, enabling dual labeling of chromatin and plasma membranes. mT/mG mice are Cre-dependent reporters that ubiquitously express membrane-targeted tdTomato before, and membrane-targeted GFP after, Cre-mediated recombination. *Thy1-YFP-H* mice express yellow fluorescent protein at high levels in motor and sensory neurons and subsets of central neurons. To induce recombination in smooth muscle cells, *αSMA*^*CreERT2*^; *tdTomato* mice were treated with tamoxifen (T5648, Sigma) at a dosage of 75 mg/kg, administered every other day for a total of three doses. All procedures with mice were approved by the Institutional Animal Care and Use Committees of National Taiwan University.

### Mouse preparation for intravital imaging

For intravital imaging of the intact ocular anterior segment, we modified a custom-designed stereotaxic mouse holder to accommodate the tilting objectives required by the vSPIM system[44]. Mice were anesthetized via Zoletil (20-40 mg/kg) and xylazine (5-10 mg/kg),. Head fixation was achieved using a three-point setup with ear bars inserted into the external auditory meatus. To provide local anesthesia, a 0.4% oxybuprocaine hydrochloride powder (O0270000, Sigma) dissolved in saline was topically applied to the corneal surface for 3 minutes. The exposed eyeball was carefully stabilized using an eye holder to avoid injury. The mouse was then positioned beneath the two tilting objectives for stable intravital imaging, with an eye gel (Vidisic Gel, Dr. Gerhard Mann Chem-pharm) was applied to the corneal surface as an immersion medium.

### Image acquisition and 3D visualization

The scan format was set to a resolution of 2048 × 2048 pixels, corresponding to a field of view of 832 µm × 832 µm. For axial resolution, the z-step size was set to 1 µm. Three-dimensional(3D) images were stitched by Imaris stitcher (version 9.3.0, Oxford Instruments) and were visualized using Imaris software (version 9.3.0, Oxford Instruments).

## Acknowledgements

We gratefully acknowledge Prof. Pierre Chambon for kindly providing the aSMA^creERT2^ mouse line. We thank Bloomington Stock Center for the fly stocks. This work was supported by National Science and Technology Council (NSTC-113-2636-B-007 -004 and NSTC-113-2311-B-007 -013 to L.A. Chu), National Taiwan University Hospital (NTUH114-E0008 to S.J. Lin), National Taiwan University (114L894201 to S.J. Lin) and Academia Sinica (AS-ASSA-114-02 to S.J. Lin), and the Brain Research Center of NTHU under the Higher Education Sprout Project funded by the Ministry of Education in Taiwan. H.C was supported by industry-university collaboration from Ministry of Education.

## Notes

### Competing Interest Statement

The authors have declared no competing interest.

