## Supplementary Information for "High-resolution volumetric intravital imaging reveals asymmetric serotonin-dependent post-shock activity in the *Drosophila* brain"

**Supplementary Fig 1**

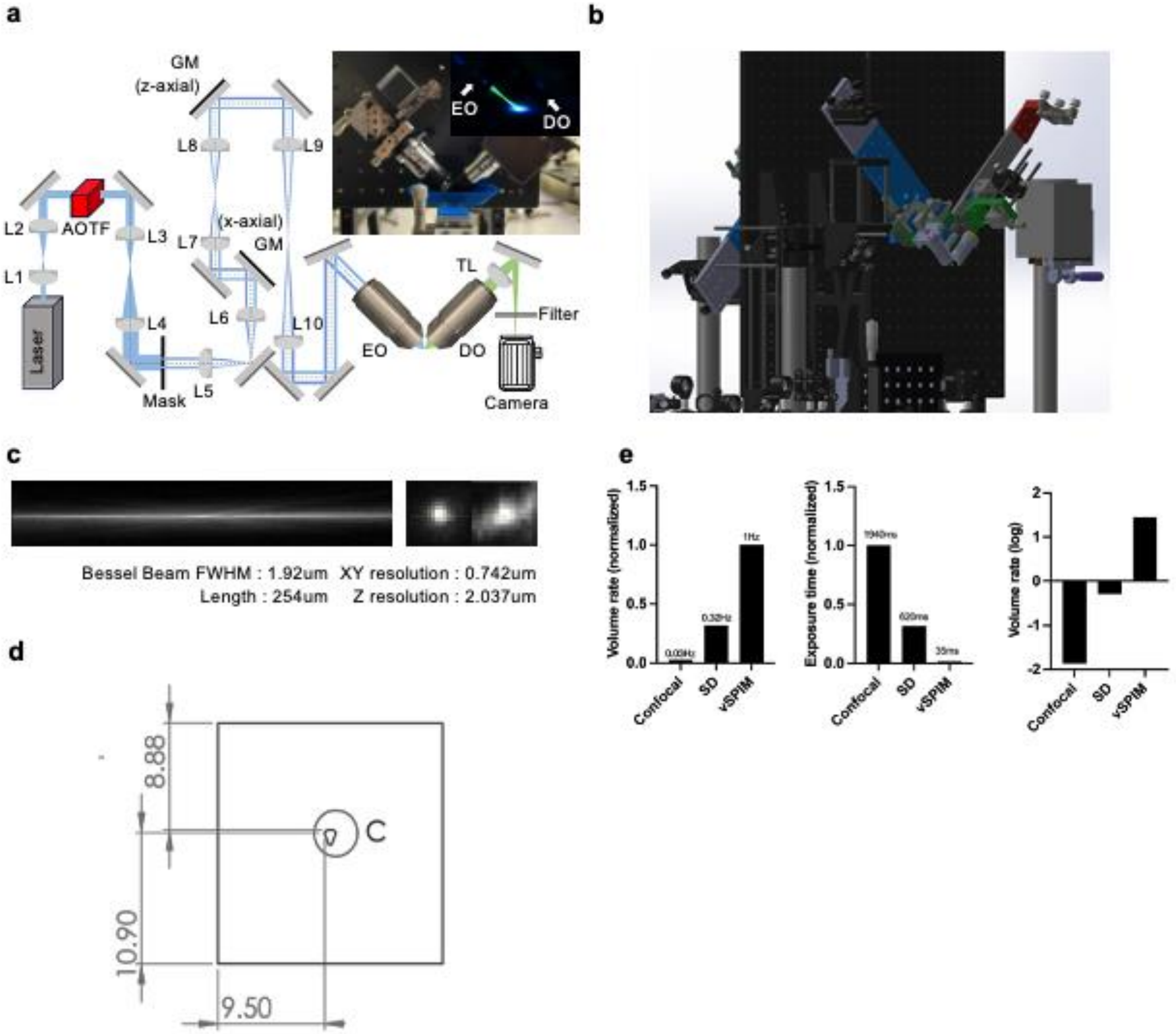


**Supplementary Figure1. Design of vSPIM and its application & system comparison.** a. Schematic diagram and photography of V-SPIM system. L1-L2, collimate lens. L3-L4, beam expander lens pair. L5-L10, relay lens. EO, excitation objective lens. DO, detection objective lens. b. System setup simulation in SolidWorks. c. Bessel beam length and thickness under detection objective. Cross-section of single fluorescence bead (200nm) showing representative point spread function (PSF) of lateral and axial resolution(lateral : 0.742$\mu m$, axial : 2.037$\mu m$). d. Customized metal plate for fly fixation(unit : mm). e. Comparison of volumetric imaging performance among vSPIM, spinning-disk confocal (SD), and confocal microscopy. Using the adult fly brain as a model (330 × 330 × 172 μm volume),. Left: normalized volume rate (V.R.) for the same exposure time (E.T.); middle: normalized exposure time required to achieve equivalent signal-to-noise ratio (SNR); right: simulated volume rate (log scale) for achieving the same SNR based on the two-prior metrics. vSPIM achieved volumetric acquisition rates 3× faster than SD and 33× faster than confocal microscopy. At equivalent signal-tobackground ratio (SNR), the minimum exposure time required for vSPIM was 35 ms, compared with 620 ms for SD and 1940 ms for confocal microscopy. Simulated volume rates at matched SNR further demonstrated >10-fold improvement over SD and nearly 10,000-fold improvement over confocal microscopy.

**Supplementary Fig 2.**

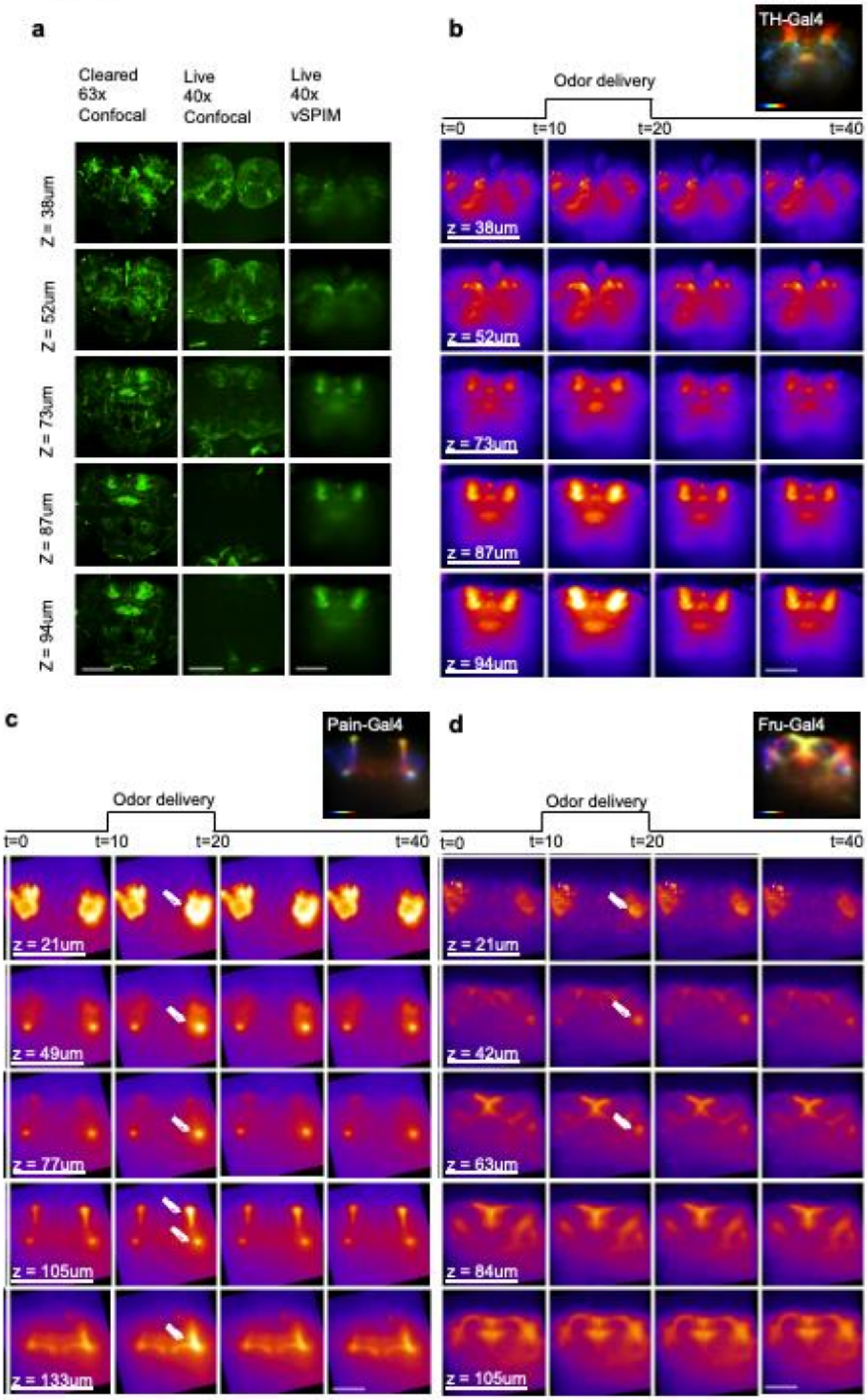


**Supplementary Figure 2. Visualization of Drosophila whole brain calcium imaging result for different neuron structure a.** Live fly brain images acquired using commercial confocal and vSPIM are compared to high-resolution images from fixed cleared fly brain. While cleared sample under confocal provides superior structural detail, vSPIM maintains higher contrast than confocal in deep regions of in-vivo fly brain imaging. **b–d**. Time-lapse volumetric imaging of odor responses in flies expressing GCaMP in different neuronal populations: b. *TH-Gal4*, c. *Pain-Gal4*, and d. *Fru-Gal4*. Each row represents dynamic calcium responses at specific depths following odor delivery .White arrows indicate regions of neural activation. (Scale bars, 100$\mu m$.)

**Supplementary Fig 3**

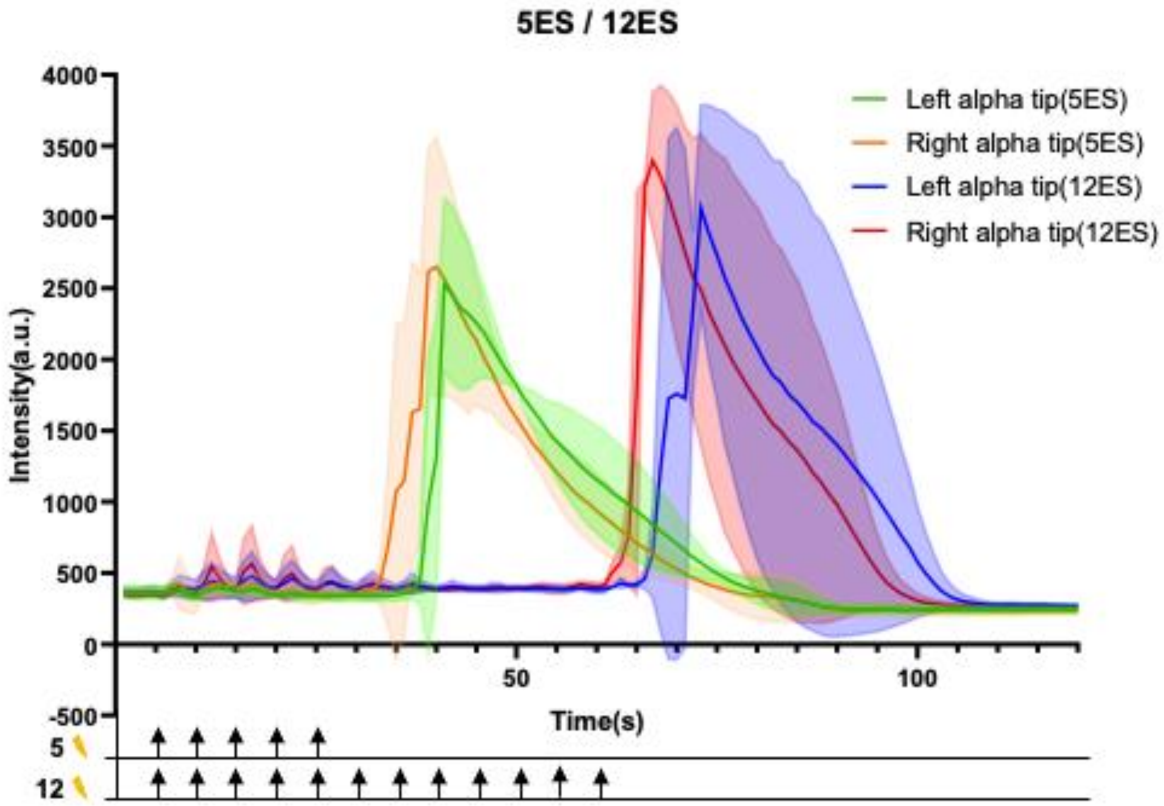


**Supplementary Figure 3. Lateralized post-shock activity dynamic in the mushroom body.** Flies with 5 times electrical shock and 12 times electrical shock intensity change profile on alpha tips at right and left brain. (5 times ES : n=3, 12 times ES : n=3).

.

**Supplementary Fig 4**

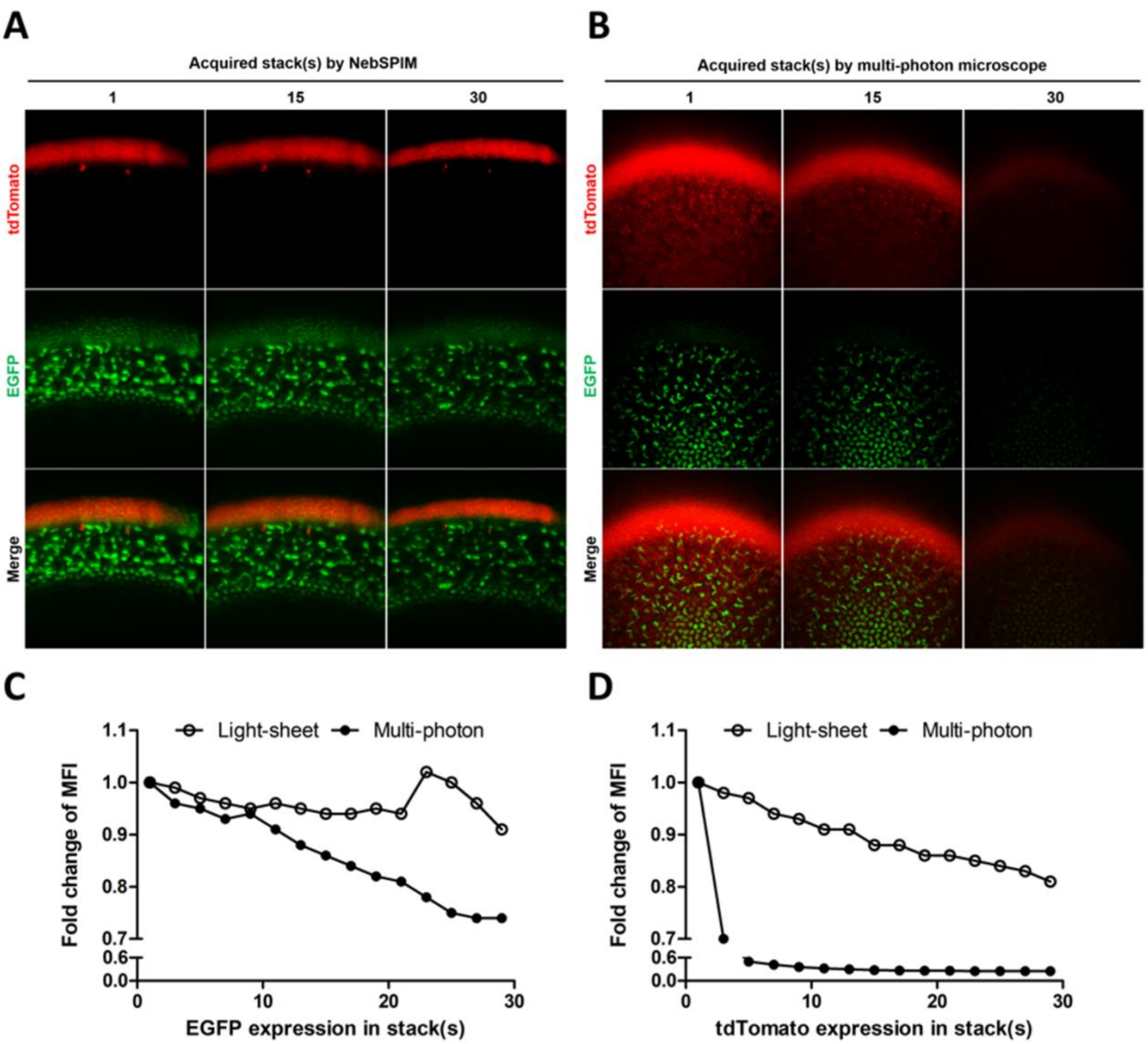


**Supplementary Figure 4. Comparison of photo-bleaching and photo-toxicity between multi-photon microscope and vSPIM.**

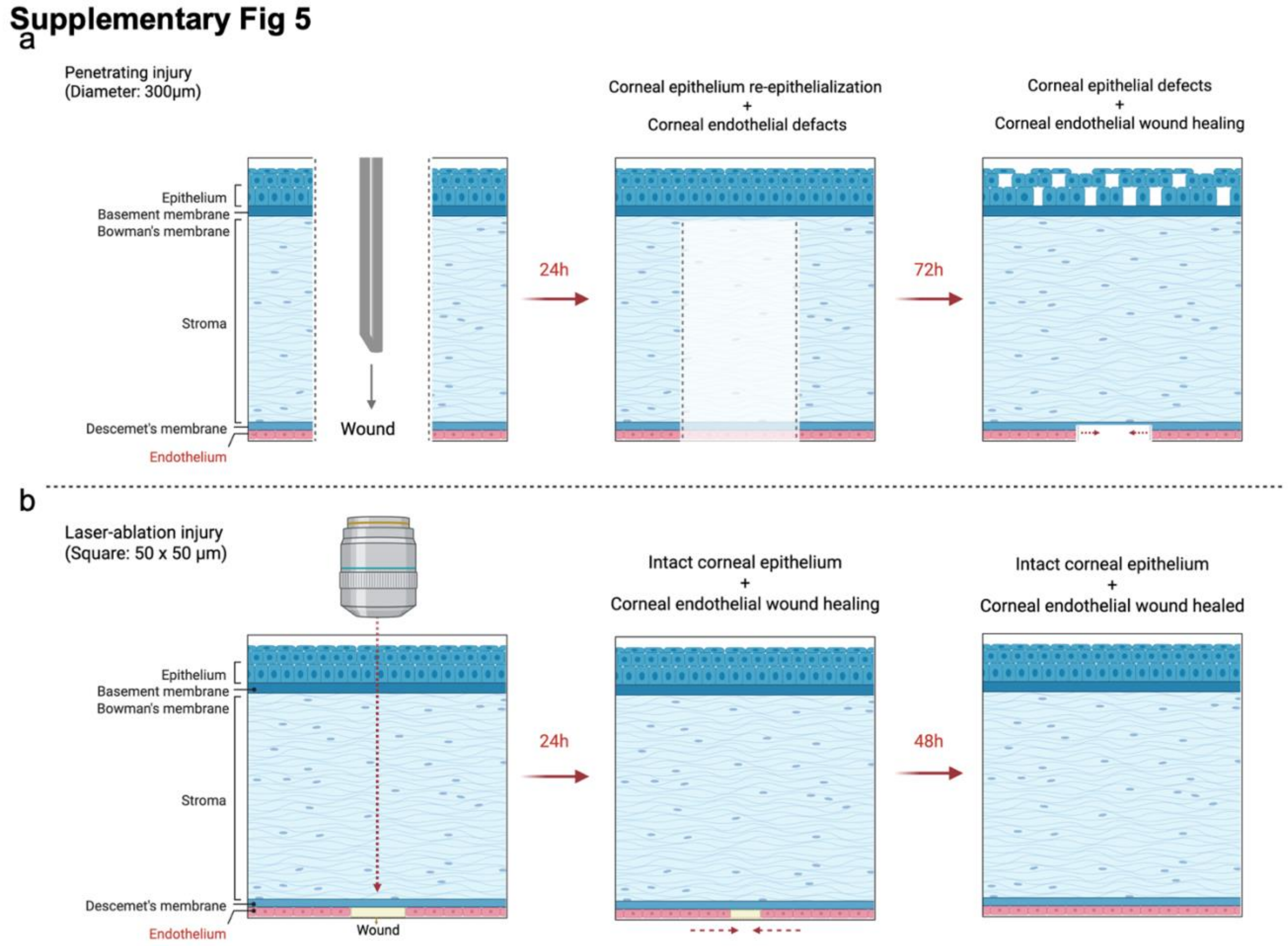


**Supplementary Figure 5. Representative diagrams of corneal injury models and subsequent healing processes. a.** Penetrating injury model. **b.** Laser ablation injury model. Schematics illustrate the distinct nature of tissue disruption and the corresponding regenerative dynamics for each model.

**Supplementary Fig 6**

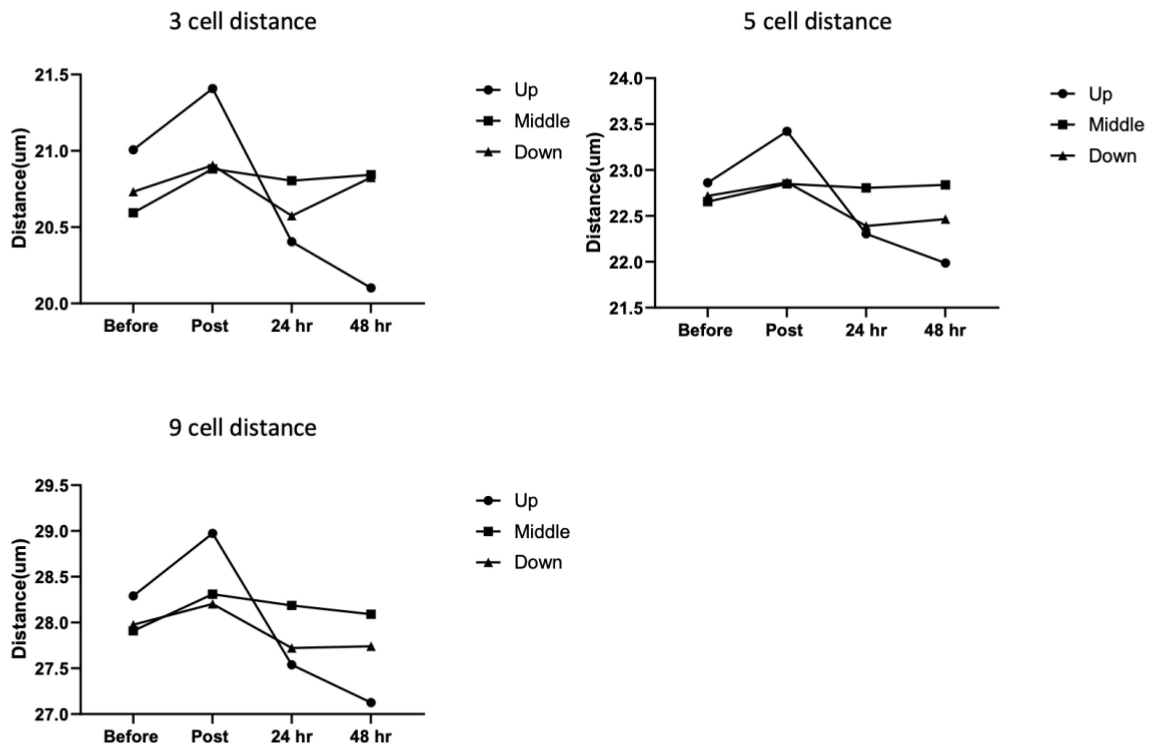


**Supplementary Figure 6. Quantification of intercellular distances in distinct regions of the corneal endothelium during wound healing.** Cell-to-cell distances were measured in the upper, middle, and lower regions of the corneal endothelium across groups of 3, 5, or 9 consecutive cells at various stages of the healing process.
